# dioscRi enables transferable prediction of clinical outcomes in multi-parameter cytometry data

**DOI:** 10.1101/2025.09.08.675022

**Authors:** Elijah Willie, Shreya Rao, Gemma Figtree, Jean Yang, Barbara Fazekas de St Groth, Helen McGuire, Ellis Patrick

## Abstract

Multi-parameter cytometry technologies enable high-dimensional analysis of immune cell populations at single-cell resolution. Deep learning has emerged as a transformative tool for analyzing these datasets, but existing methods often struggle with transferability across datasets due to technical variability, batch effects and identification of biologically relevant cell populations, limiting their utility in clinical research. We present dioscRi, a transferable deep learning framework that integrates a maximum mean discrepancy variational autoencoder for normalization and de-noising, enhancing cross-dataset compatibility. Changes in cell type proportions and marker expression are identified by structuring these features within biologically or empirically derived cell type hierarchies. These hierarchies are incorporated directly into an overlapping group LASSO model, improving the prediction of clinical outcomes. When applied to a coronary artery disease study, dioscRi recapitulated several known immune associations. Benchmarking across multiple datasets demonstrated dioscRi’s ability to generalize and outperform existing methods, establishing it as a versatile and interpretable tool for cytometry data analysis.

## Introduction

Multi-parameter flow and mass cytometry technologies enable high-dimensional profiling of millions of cells in a single patient sample, a capability that can support a range of clinical applications including diagnostics, prognostics, and therapeutic monitoring [1–3]. These platforms can measure dozens of markers across millions of cells per sample, providing deep insights into immune states, tumor microenvironments, and signaling dynamics across diseases [4, 5]. However, translating these insights into clinical practice requires computational methods that can scale to large datasets, remain stable in the presence of technical variability, and generalize across cohorts without retraining. To be clinically actionable, such methods must also produce outputs that are interpretable to researchers and clinicians.

Deep learning encompasses a range of methods that have shown promise for cytometry analysis, owing to their ability to model the complexity of cellular systems. Architectures such as autoencoders (AE), variational autoencoders (VAE), and convolutional neural networks (CNN) are commonly used to extract meaningful representations from highdimensional data while remaining relatively robust to noise and variability [6]. For example, MoE-Sim-VAE [7] combines VAEs with Gaussian mixture models to enable unsupervised identification of cell types, implicitly separating biological variation from technical artifacts. DGCyTOF [8] and DeepCNN [9] apply autoencoders for supervised cell-type classification, with DeepCNN incorporating a calibration step to address batch effects across datasets. However, such calibration requires prior knowledge of batch structure, limiting its applicability in clinical settings where classification of a sample should be done independently of other samples. For patient classification, models such as CellCNN [10] and DeepCNN use CNNs to learn complex relationships directly from single-cell measurements. Recognizing the importance of interpretability, DeepCNN also employs post-hoc permutation tests and decision trees to identify marker-level associations. Because these interpretability steps are applied after model training, they do not constrain the model during learning and may not reflect the true decision boundaries. As a result, existing approaches are not designed to generalize to unseen datasets and typically lack mechanisms for structured interpretation, making them difficult to apply confidently across cohorts or translate into clinical insight.

A longstanding solution to the challenges of interpretability and technical variability in cytometry has been the use of cell-type hierarchies [11, 12]. These hierarchies are typically constructed through manual gating, a conventional technique in fluorescence and mass cytometry that defines cell populations by drawing boundaries on a series of two-dimensional marker expression plots which, taken together, encompass a complex phenotype including expression of multiple markers. Analysts begin by identifying broad immune categories, such as CD4 and CD8 T cells, NK cells and B cells, and progressively refine these into more specific subsets such as regulatory versus conventional CD4 T cells, naïve versus memory conventional CD4 T cells and so on. This hierarchical structure not only facilitates interpretation but also provides robustness to technical variation, as comparisons between child and parent populations help normalize differences across samples. Manual gating remains widely used because it reliably identifies cell subpopulations with known biological functions while effectively accounting for technical variability. However, it requires considerable expertise, and is very time-consuming, especially when sample-to-sample variation from biological or technical sources requires that multiple gates be adjusted for each sample. Hence the search for computational methods that can replicate and extend the complex cell subset structures within data derived from increasingly high-plex technologies [13, 14]. Recent approaches aim to connect cell populations to biological or clinical outcomes by leveraging hierarchical relationships in ways that preserve interpretability and improve scalability [15, 16] Embedding hierarchical relationships into deep learning models could improve their resilience to technical variation while preserving biologically meaningful structure.

To address the need for models that are both interpretable and transferable across batches and cohorts, we present dioscRi, a deep learning framework for predicting clinical outcomes from multi-parameter cytometry data. DioscRi combines a Maximum Mean Discrepancy variational autoencoder (MMD-VAE) to learn a transferable normalization scheme, with an overlapping group LASSO model that incorporates biologically or empirically derived cell-type hierarchies. This design enables the model to generalize across datasets without requiring batch labels, while producing structured outputs that reflect known immune relationships. We demonstrate the utility of dioscRi through its application to a coronary artery disease cohort, where it recapitulates a subset of previously published immune associations [17]. We further benchmark its performance across multiple public datasets, showing that dioscRi consistently outperforms existing deep learning approaches. Together, these results establish dioscRi as a robust and generalizable framework for clinically focused cytometry analysis.

## Methods

### Overview of dioscRi

dioscRi consists of various steps: (i) Transferable data normalization and denoising using a Maximum Mean Discrepancy Variational Autoencoder (MMD-VAE), (ii) Generation of cell types through unsupervised clustering, (iii) Extraction of cell type and sample-level features, (iv) Hierarchical grouping of cell type-level features, (v) Clinical outcome prediction using overlap group lasso, and (vi) Model visualization and interpretation.

### Data Pre-processing

For all datasets, we applied an *arcsinh* transform (*y* = *arcsinh*(*x/*5)). Following transformation, each data set was sub-sampled to 10,000 cells from each sample and combined for analysis.

Before training the MMD-VAE model, a Min-Max scaling to (0, 1) was applied.

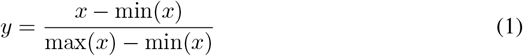

Min-max scaling is commonly employed to improve the training efficiency and convergence of deep learning models.

### Maximum Mean Discrepancy Variation Auto Encoder Architecture

The dioscRi architecture integrates an encoder and decoder within a Variational Autoencoder (VAE) framework. It is enhanced by the Maximum Mean Discrepancy (MMD) to better match the latent space distribution to the prior, improving adaptability to non-Gaussian data distributions [18, 19]. The Maximum Mean Discrepancy (MMD) penalty was implemented to enforce distributional similarity between the encoded latent variables and a prior distribution, following the method described in [20]. An unbiased U-statistic estimator of the MMD was computed using an Inverse Multi-Quadratic (IMQ) kernel. We summed IMQ kernels computed at varying scales to capture multi-scale relationships. The encoder compresses the input data into a lower-dimensional representation, thereby retaining only the most pertinent information by filtering out noise. The decoder then reconstructs this encoded data to its original form by minimizing the difference between original input and the decoded output.

The model can be formulated as:

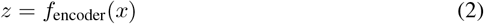

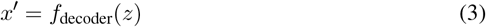

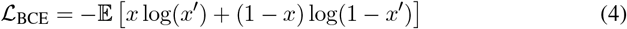

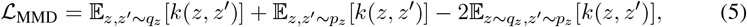

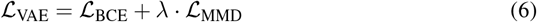

Here, *f*_encoder_ is the encoder function that compresses the input data *x* into a lower-dimensional latent representation *z*. The decoder function *f*_decoder_ reconstructs the input data from *z* back to its original form *x*^*′*^. The Binary Cross Entropy loss ℒ_BCE_ measures the difference between the input and the reconstructed output. For the MMD loss, *k*(·, ·) is the IMQ kernel. This penalty encourages the encoded latent representation to align with the desired prior distribution while allowing flexibility for modeling complex data distributions. The overall loss ℒ_VAE_ combines the Binary Cross-Entropy loss ℒ_BCE_ with the MMD regularization term ℒ_MMD_, balanced by a hyper-parameter *λ* that controls the influence of the MMD term.

### MMD-VAE Implementation

The model architecture includes an encoder that reduces the dimensionality of the input data through dense layers with ReLU and GELU activations, followed by a decoder that reconstructs the input using sigmoid activation. The training objective optimizes a total loss, combining binary cross-entropy for reconstruction and Maximum Mean Discrepancy (MMD) to align the latent space distribution with a standard normal prior.

The MMD-VAE model is implemented in R using the Keras library [21], with RMSprop [22] as the optimizer and a default learning rate of 0.001. The MMD-VAE is trained for 100 epochs with a batch size of 32, and the latent dimension is conservatively set to balance information retention. The model is validated using a separate dataset during training. The MMD-VAE comprises three hidden layers with layer sizes differing for each data depending on the number of features. Table 2 lists the hyper-parameters used for each dataset.

**Table 1.** Benchmark datasets. Five published datasets were used to benchmark and compare the performance of dioscRi to other deep-learning models. “Name” refers to each dataset throughout the manuscript

| Name | Technology | Description | Num Samples | Clinical Outcome |
| --- | --- | --- | --- | --- |
| Wagner et al Breast Cancer | CyTOF | Predicting tumor in breast cancer samples | 168 | Tumor/non-tumor |
| CMV - Study SDY519 | CyTOF | Predicting positive vs negative CMV titer results in influenza samples | 472 | Positive/negative |
| Bioheart-CT Discovery | Mass cytometry | Discovery study for BioHEART-CT | 111 | Gensini_bin = 1/0 |
| Bioheart-CT Validation | Mass cytometry | Validation study for BioHEART-CT | 58 | Gensini_bin = 1/0 |
| Matthew et al COVID19 | Flow cytometry | Profile of CD8+ Non-Naïve T Cells to distinguish recovered from COVID-19 vs. healthy | 214 | COVID-19 recovered / healthy |

**Table 2.**
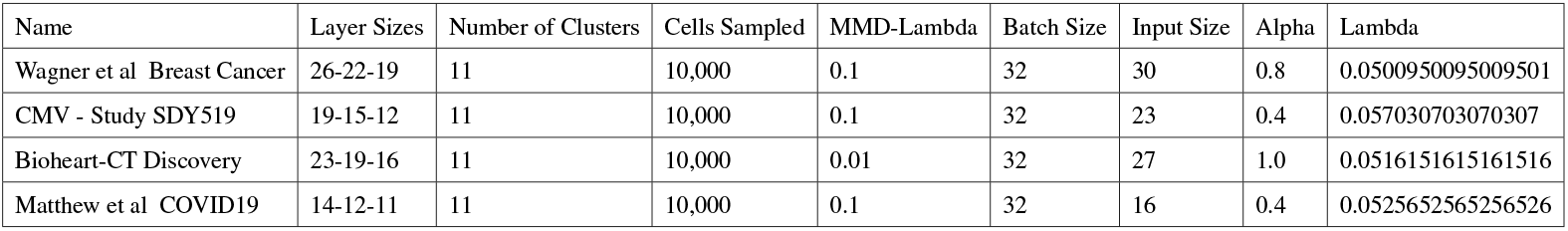
Hyper-parameters used for each dataset when fitting dioscRi MMD-VAE normalization and Overlap-Group Lasso.

### Selection of Representative Sample for MMD-VAE Training

To reduce the total size of the training data while retaining core biological information, we select representative samples for training the MMD-VAE model. We ensure the model captures key biological variations without redundancy by selecting samples with the most representative characteristics. To achieve this, we use the method described in [23], where each sample is evaluated based on its average covariance Frobenius norm between it and other samples.

The Frobenius norm quantifies the difference between the covariance matrices of pairs of samples, effectively measuring how similar or different each sample is to others. Formally, the Frobenius norm between samples *i* and *j* is defined as:

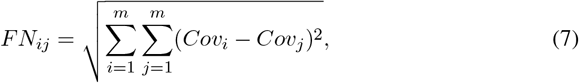

where *Cov*_*i*_ and *Cov*_*j*_ are the covariance matrices for samples *i* and *j* respectively. The stable sample is selected by returning the sample that satisfies:

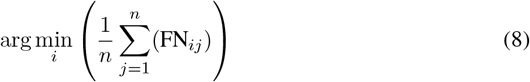

Using this criterion, we select the top *K* samples with the minimum average Frobenius norms as the set of reference samples, with *K* set to 2 in all experiments.

### Unsupervised Clustering and Cell Type Classification

To generate cell types from the normalized training data, we use the FuseSOM algorithm [24], which combines multiple similarity metrics through multiview ensemble learning and hierarchical clustering to create cell types.

For the testing data, cell types were generated by training a classifier on the cell types identified in the training data. A Linear Discriminant Analysis (LDA) model was used for all cell type classifications. The embeddings from the VAE model served as features for unsupervised clustering and manual gating. Cell type classification models were fit using the *Caret* R package.

### Feature Engineering

After cell type identification, we generate a set of features, including the logit of the cluster proportions in each sample and the mean marker expression per cell type in each sample.

The logit proportion for a cluster *k* in sample *i* is defined as:

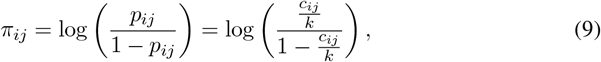

where *n* is the number of samples, *k* is the number of cells in each sample, *m* is the number of clusters, and *c*_*ij*_ is the count of cells in sample *i* for *i* = 1, 2, …, *n* that belong to cluster *j* for *j* = 1, 2,, *m*.

The mean marker expression per cell type across all samples is defined as:

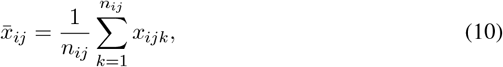

where *x*_*ijk*_ is the marker expression value of the *k*^*th*^ cell belonging to cell-type *j* and sample *i*. The value*n*_*ijk*_ is the total number of cells of belonging to cell-type *j* in sample *i*.

### Incorporating empirically or biologically informed grouping structure

Before classifying samples, we integrate cell type hierarchies into the proportions, using either biologically informed or empirically derived groupings. To empirically derive groupings we construct a hierarchical clustering tree based on Pearson correlation distance and Ward linkage, generating all possible groupings by cutting the tree at various heights. Alternatively, biologically derived hierarchies are predefined using prior knowledge of cell-type relationships by respective domain experts. These hierarchies, whether empirically or biologically informed, are combined with the marker means for each cell type to create the final set of features used for classification.

### Group Logistic Lasso with Overlapping Groups

The Lasso (Least Absolute Shrinkage and Selection Operator) is a regularization method used for feature selection and model regularization [25]. Given a matrix *X* of predictors with *n* samples and *p* features, and a binary response vector *y*, the Lasso minimizes the following objective function:

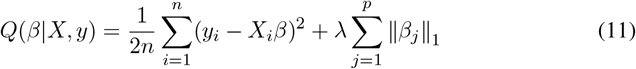

where *β* represents the coefficient vector, 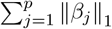 is the *ℓ*_1_-norm penalty, and *λ* is a regularization parameter that controls the level of sparsity in the solution. By penalizing the absolute values of the coefficients, Lasso encourages sparsity, driving some coefficients to zero and excluding irrelevant features. This approach is particularly effective in high-dimensional settings, where it reduces overfitting and yields interpretable models.

The Group Lasso extends this framework by promoting sparsity at the group level, enabling feature selection based on predefined groups of predictors [26]. Assume that the predictors are partitioned into *J* non-overlapping groups *G*_1_, *G*_2_, …, *G*_*J*_, with each group containing a subset of features. The Group Lasso objective is:

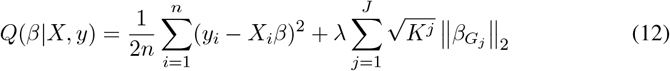

where 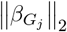 denotes the *ℓ*_2_-norm of the coefficients within group *G*_*j*_, and *K* is the number of elements in group j.

We adapt the Group Lasso with a logistic loss function for binary outcomes to handle classification. Given a binary response vector *y* with entries *y*_*i*_ ∈ {0, 1}, the logistic loss is defined as:

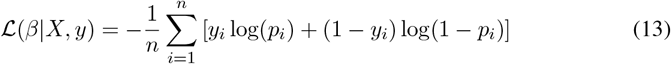

where

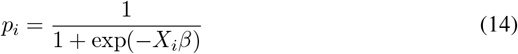

represents the probability of *y*_*i*_ = 1 given *X*_*i*_. The Group Lasso with logistic loss thus becomes:

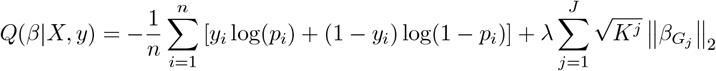

This formulation encourages sparsity in the group structure, making it well-suited for binary classification tasks involving predefined groups of features.

The Overlapping Group Lasso further generalizes this approach by allowing predictors to belong to multiple groups [27]. This extension is particularly useful in cases where variables naturally fall into overlapping categories, such as biological pathways where genes may belong to multiple pathways, or immune cell features that are shared across related cell types within a hierarchy. To accommodate overlapping groups, we introduce a latent structure by decomposing the coefficient vector *β* into group-specific latent vectors *γ*_*j*_, each associated with a group *G*_*j*_. Let 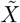 be the expanded design matrix with duplicated columns for overlapping variables. The objective for the Overlapping Group Lasso is:

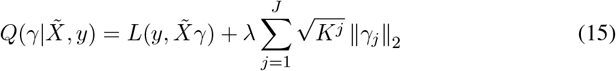

where 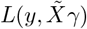 denotes the logistic loss, *γ* = (*γ*_1_, …, *γ*_*J*_) represents the concatenated vector of group-specific coefficients, and ||*γ*_*j*_||_2_ applies the *ℓ*_2_-norm penalty within each group. The overlapping group structure enables shared predictors to have multiple group-specific effects, enhancing model flexibility. After fitting the model, the original coefficients *β* are reconstructed as:

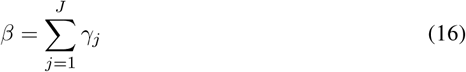

To further regularize overlapping cases, we add a ridge penalty term, defining the final objective as:

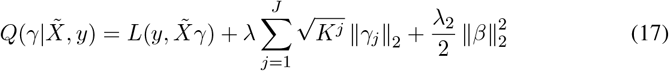

where *λ*_1_ = *αλ* and *λ*_2_ = (1 − *α*)*λ*. To select the optimal *α* and *λ*, we fit models with different *α* values over the range [0,1], *α* values over the range [0.05, 1] and compute the deviance criterion for each model. The best *α* and *λ* values are chosen using the elbow method applied to the deviances. We implemented the overlapping group lasso using the grpregOverlap R package [27], modified to support logistic loss for binary classification. This approach provides a flexible and interpretable solution for feature selection when variables belong to multiple groups.

In the overlapping group lasso model, each coefficient vector *γ*_*j*_ represents the contribution of a specific group *G*_*j*_ to the overall model, allowing for a unique interpretation of overlapping features. Since 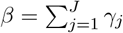, the original coefficient vector *β* is constructed by summing the group-specific latent effects *γ*_*j*_. This decomposition means that each predictor can be influenced by multiple groups simultaneously, with each *γ*_*j*_ capturing the effect of group *G*_*j*_ independently. Consequently, the magnitude and direction of *γ*_*j*_ can be interpreted as the specific impact of group *G*_*j*_ on the predictor, highlighting how particular groupings contribute to the model’s predictive power. When interpreting the model, non-zero *γ*_*j*_ values indicate that the group *G*_*j*_ plays a significant role, while the summed *γ* coefficients in *β* provide the overall effect of each predictor. This approach improves interpretability by revealing which features are selected and which groups drive their inclusion, offering insights into overlapping pathways or shared structures across groups.

### Implementation of other approaches for benchmarking comparisons

#### Training of cyCombine and iMUBAC

We benchmarked the performance of the normalization performance of dioscRi against other normalization methods that explicitly consider batch information. These methods included cyCombine [28] and iMUBAC [29]. For analysis, both cyCombine and iMUBAC were run with default parameters.

#### Training of CellCNN and DeepCNN

The predictive performance of dioscRi was benchmarked against CellCNN [10] and DeepCNN [9]. Since both methods work directly on FCS files, after splitting each dataset into training and testing, the resulting cells were written back to FCS files for predictive modeling using these methods.

CellCNN was run using an optimal set of hyper-parameters (filter number = 5, *l*_2_ coefficient = 1*e* − 4). The Adam optimizer was used for loss optimization, with a learning rate of 0.01 over 500 epochs. After training, the model was evaluated using the testing set.

DeepCNN was run using the optimal set of hyper-parameters used in the original paper (number of filters = 3, dropout rate equals 0.2). The Adam optimizer was used for loss optimization, with a learning rate of 0.01 over 500 epochs. After training, the model is evaluated using the testing set.

### Model Evaluation

Various evaluation metrics were used to evaluate components of dioscRi.

### Evaluation of Normalization by Clustering

After data normalization, the Adjusted Rand Index (ARI) was used to assess clustering performance. The ARI metric assesses the overlap between two sets and takes on values ranging from 0 to 1, with 0 indicating no overlap and 1 indicating perfect overlap. The ARI score can be defined as:

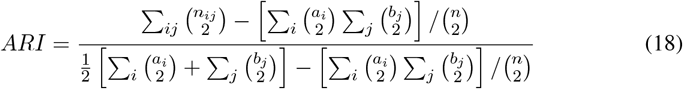

where *n*_*ij*_ is the number of pairs of elements that are in the same set in both clusterings, *a*_*i*_ is the total number of pairs in the same set for the first clustering, *b*_*j*_ is the total number of pairs in the same set for the second clustering, and *n* is the total number of elements.

### Evaluation of Outcome Prediction

The ROC-AUC metric was used to assess the performance of dioscRi and competing methods for predicting clinical outcomes. The metrics are defined using the true positive (TP), true negative (TN), false positive (FP), and false negative (FN) values.

The ROC-AUC measures the area under the Receiver Operating Characteristic (ROC) curve, which plots the true positive rate (TPR) against the false positive rate (FPR) at various threshold settings. The AUC score is a single scalar that summarizes the performance across all possible classification thresholds, with 1 indicating perfect classification and a value of 0.5 indicating no better than chance. The formula for TPR and FPR are:

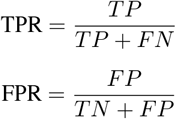

### Datasets

We employed a variety of datasets across different technologies and diseases. Brief descriptions of each dataset are provided below, with more details outlined in Table 1.

### BioHEART-CT

To evaluate the dioscRi framework, we used mass cytometry data generated from the BioHEART-CT study [17, 30]. This prospective longitudinal cohort study to identify immune cell populations associated with coronary artery disease (CAD). Cell profiling was performed using mass cytometry with 41 tagged antibodies, enabling a detailed analysis of cellular characteristics. The major clinical outcome used in the previously published analysis [17] was the Gensini score, a continuous measure of coronary artery disease (CAD) severity. For prediction tasks, we binarized this outcome: samples with a Gensini score of 0 were labeled as CAD−, and those with a score greater than 0 were labeled as CAD+. The original study [17] included discovery and validation cohorts that underwent expert manual gating to identify more than 100 subpopulations. The discovery cohort of 111 samples across 6 batches comprising a total of 41,350,484 cells was then analyzed using a logistic regression model to identify 18 subpopulations whose size was identified as most different between CAD+ and CAD−. The sizes of the same 18 subpopulations were manually gated in the validation cohort consisting of 58 samples across 3 batches. Importantly, the validation cohort was acquired using a mass cytometry platform that introduced significant time-dependent changes in multiple signals [31], so the expert gating included preliminary time gates to exclude artifactual data. For the current study, the discovery cohort was downsampled to include 1,106, 443 cells. The validation cohort was downsampled to 580,000 cells without applying any time gates. For the discovery cohort, 11 manually gated cell populations, identified as detailed in Supplementary Figure 4 of the original manually gated analysis [17] were used as the basis for cell type prediction. The 11 populations were conventional CD4+ T cells (Tconv, mean 38%), CD4+ regulatory T cells (Treg, mean 1.9%), CD8+ T cells (CD8hi mean 17%), CD8lo-neg T cells (CD8lo mean 6.2%), B cells (mean 10%), NK cells (mean 10.2%), CD14+ monocytes (mean 12%), CD16+ monocytes (mean 2.1%), plasmacytoid dendritic cells (pDC mean 0.36%), CD1c+ dendritic cells (CD1c DC mean 0.65%) and CD141+ dendritic cells (CD141 DC mean 0.044%). Of the 41 metal-tagged antibodies used to assess marker expression in the original mass cytometry data, 27 were included as markers in the discovery and validation datasets.

### Breast cancer tumor

This dataset comprises 144 human breast tumor samples and 50 non-tumor tissue samples measured with mass cytometry[32]. The primary objective was to characterize the features of breast cancer ecosystems and their correlations with clinical data. The study evaluated 73 proteins in approximately 26 million cells, utilizing tumor-focused and immune cellcentric antibody panels. This extensive profiling helps understand the protein expression patterns and cellular interactions within breast cancer and adjacent non-cancerous tissues. We analyzed 194 samples, predicting each as tumor vs. non-tumor. The dataset was split into 70% training and 30% testing.

### Cytomegalovirus

This dataset encompasses data from nine human immunology studies (SDY112, SDY113, SDY305, SDY311, SDY315, SDY472, SDY478, SDY515, and SDY519) featuring 596 peripheral blood mononuclear cell (PBMC) samples from 313 subjects across 472 samples [33–36]. Diagnosing latent Cytomegalovirus (CMV) is challenging since it is usually asymptomatic with limited observable effect on the immune system. These studies aimed to identify immune responses to latent CMV infections. Our analysis of this dataset followed that used in the DeepCNN manuscript [9], which used studies SDY519 and SDY515 as testing and validating datasets and the rest for model training. Clinical response is individuals having a positive or negative cytomegalovirus (CMV) infection. A set of 23 markers that were common to all nine studies.

### COVID-19 PBMC CD8+ non-naive T cells

This dataset is derived from a high-dimensional flow mass cytometry analysis of COVID19 samples, alongside comparisons with recovered individuals and healthy controls [37]. The study involved an integrated analysis of approximately 200 immune features. These immunological data were combined with about 50 clinical features, providing a comprehensive overview that facilitates understanding of the relationships between the immunology of SARS-CoV-2 infection and various clinical outcomes, including disease patterns, severity, and progression. We restricted analysis to CD8+ non-naive T cells and used COVID-19 recovery as the clinical response of interest. The dataset was randomly split into 70% training and 30% testing sets.

## Data Availability

All datasets used in this study are publicly available. Data were downloaded following the instructions of their respective publications.

## Results

### Introducing dioscRI

Here we present dioscRi, an interpretable framework for analyzing multi-parameter cytometry data. This framework integrates an MMD-VAE with a hierarchical group LASSO tailored for the prediction of clinical outcomes (Figure 1). The input to the framework consists of raw gene or protein (markers) expression data of all samples, where rows represent individual cells and columns correspond to markers. The model output can include normalized expression data, unsupervised cell clusters, sample-level clinical outcome prediction, and the associations of cell-type proportions and/or markers with clinical outcomes. The MMDVAE provides a transferable normalization scheme, ensuring it can be applied to unseen data. The overlapping group LASSO uses a biologically or empirically derived grouping of cell-type proportions and marker means per cell-type to accurately predict clinical outcomes and uncover associations. The model outputs a tree structure that can assess the associations of these proportions and marker means within cell-types with predictions. In addition, the framework allows for the inclusion of additional clinical information, such as age and gender, which can enhance the precision of clinical outcome predictions.

**Figure 1.**
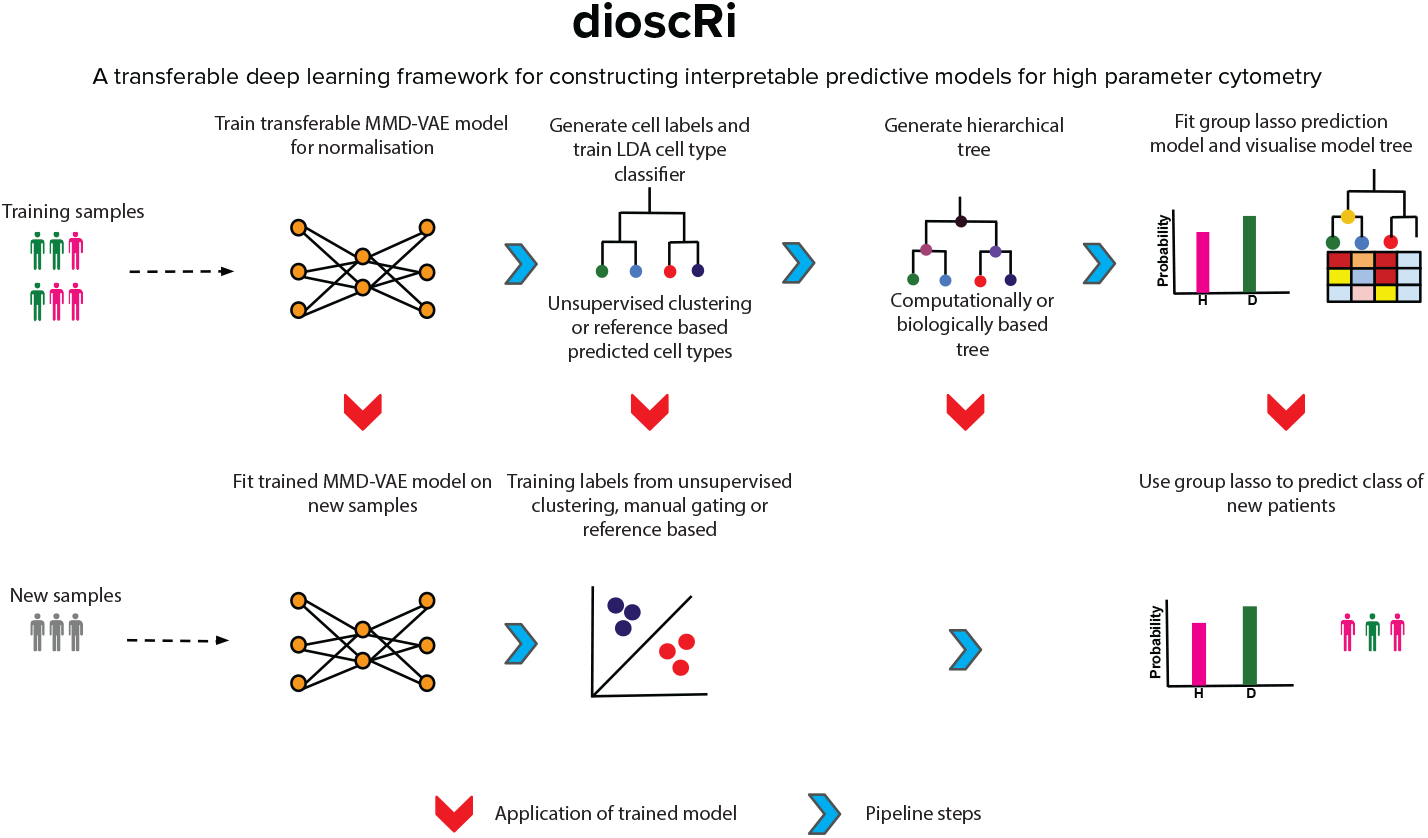
Overview of dioscRi: this new framework uses: (I) a robust MMD-VAE to train a normalization filter scheme that can applied to new data; (ii) linear discriminant analysis trained using either manually gated cell populations or unsupervised clustering to classify cell types; (iii) a hierarchical group LASSO that uses either a biologically-defined or computationally derived tree to combine predicted cell type or cluster proportions and marker means in each sample to predict clinical outcomes; and (iv) a visualization of the associations between cluster or predicted cell type features and clinical predictions using heatmaps and the tree structures.

### dioscRi Effectively Normalizes Unseen Samples

To assess the effectiveness of dioscRi’s normalization, we applied it to a mass cytometry study of coronary artery disease and evaluated its ability to reduce technical variability across samples. A discovery cohort of 111 samples was used to train the normalization Variational Autoencoder, which was then applied to a validation cohort of 58 samples. Once trained, the model was applied consistently across both cohorts to ensure a unified normalization strategy. The impact of normalization was first evaluated by examining co-expression patterns using biaxial scatter plots of CD3 and HLA-DR for samples 87 and 88 (Figures 2A–D). In the raw data, broad and overlapping distributions indicated sample-to-sample inconsistency. Following normalization, these distributions became more compact and well separated, supporting improved consistency in population structure and enhancing comparability across samples for downstream analyses. Next, density plots of CD3 (Figures 2E and 2F) revealed substantial improvement following normalization. In the raw data (Figure 2E), distributions varied notably between samples suggesting technical variability. After applying dioscRi (Figure 2F), marker distributions became more closely aligned, reducing non-biological variation and preserving meaningful signal.

**Figure 2.**
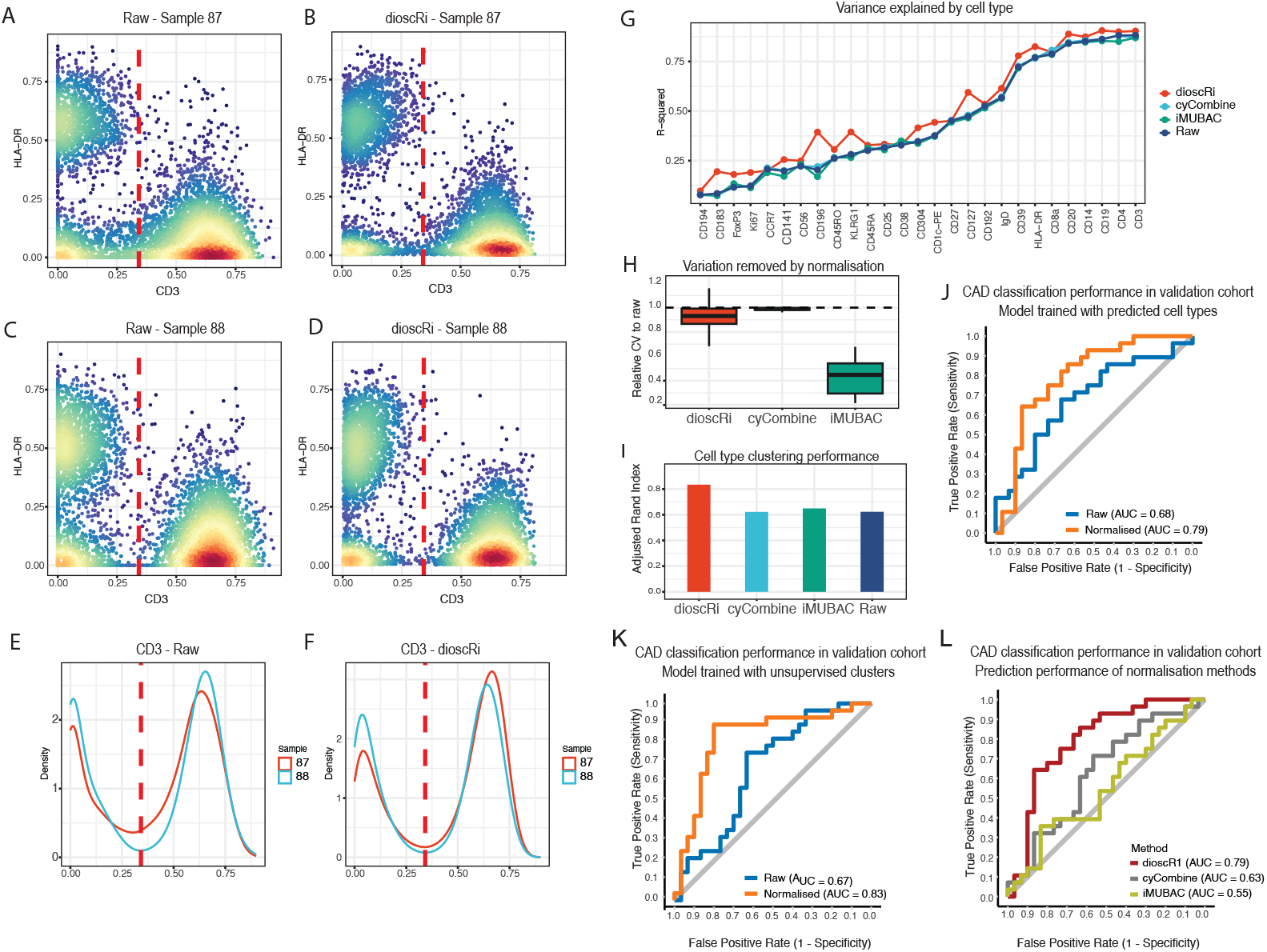
(A-D) Biaxial scatter and (E-F) density plots illustrate the effect of normalization on marker expression and gating consistency in samples 87 (validation cohort) and 88 (discovery). In the raw data (A, C, E), marker distributions are broad, clusters are poorly defined, and CD3 alignment is inconsistent, particularly in the validation cohort sample. After dioscRi normalization (B, D, F), distributions become tighter, cell populations separate more clearly, and CD3 expression aligns across samples. Red dashed lines indicate gating regions for specific subsets, highlighting reduced variability and preservation of biologically relevant features post-normalization. (G) R2 values from the relationship between predicted cell types and each marker for three normalization strategies and the raw data. (H) Coefficients of variation for each marker before and after normalization. (I) Adjusted Rand Index is used to evaluate if clustering on the different normalized datasets recovers populations similar to the manually gated cell types. Finally, AUC of models classifying CAD using dioscRi improves patient classification in (J) models trained using cell types predicted from manually gating and (K) models trained with clustered cell populations. (L) Comparison of CAD classification reveals a higher AUC for dioscRi normalization compared with cyCombine or iMUBAC normalization.

### dioscRi Normalization Reduces Variance and Improves Cell Annotation

We next assessed the effect of normalization across all markers. We first computed the *R*^2^ from linear models of marker expression against predicted cell type (Figure 2G). DioscRi consistently produced the highest *R*^2^ values relative to raw data and other methods, indicating effective noise reduction while preserving biological signal. Next, comparisons of relative coefficient of variation (CV) (Figure 2H) showed that dioscRi modestly reduced variability (median = 0.92) while maintaining a tight distribution across markers; cyCombine showed no change (median = 1.00) and iMUBAC achieved the largest reduction (median = 0.45), which when interpreted in the context of the *R*^2^ results suggest this is driven by loss of meaningful biological heterogeneity. We then confirmed this balance at a more granular level by performing marker-level analysis of variability and signal intensity across samples (Supplementary Figure S1). In the raw data, markers such as CD3, CD4, and HLA-DR exhibit higher variance, consistent with both biological reality and potential technical artifacts. In combination, these results indicate that dioscRi removes sufficient technical noise to stabilize CV without over-compressing true variation.

To assess the impact of normalization on cell type identification, we then performed clustering on the data that had been normalized by the different approaches. Models were also trained with either unnormalized or normalized data using the 11 manually gated cell populations in the discovery cohort to predict cell types in the discovery and validation cohorts. We assessed the concordance of the clustering and predicted cell types using the adjusted Rand index (ARI; Figure 2I). DioscRi achieved an ARI of 0.83, outperforming cyCombine (ARI = 0.62), iMUBAC (ARI = 0.65), and the raw data (ARI = 0.62). The Rand index value of 0.83 for dioscRi indicates that the gated and clustered cell types would have biologically significant differences. However, in comparison to cyCombine and iMUBAC, the results underscore dioscRi’s capacity to preserve biologically meaningful structure even without explicit batch labels by producing clustering results that align with known cell-type assignments.

### dioscRi Normalization Improves Patient Classification

To evaluate the impact of normalization on predictive performance, we applied dioscRi to a coronary artery disease dataset and compared outcomes using normalized and unnormalized data. Classifiers to predict coronary artery disease (CAD) were constructed that used predicted cell type proportions and marker means as input features, as well as the biologically-informed hierarchy of the manually gated cell populations. After training models on the discovery cohort, normalization led to substantial improvements in predictive performance with AUC in the validation cohort improving from 0.68 to 0.79 (Figure 2J). To demonstrate that the normalization performance benefits of dioscRi are not solely dependent on expertly derived hierarchical relationships, we then derived 11 cell populations from unsupervised clustering with FuseSOM [24] and an empirical hierarchical tree as described in the methods. For this clustering-based model, normalization increased the AUC from 0.67 to 0.83 (Figure 2K). Finally, we compared the benefit of dioscRi’s transferable normalization with the two established batch correction methods, cyCombine and iMUBAC. Both cyCombine and iMUBAC used information from the discovery and validation sets for normalization, while dioscRi only used the discovery data to train its normalization. Following normalization with each method, the resulting data were processed through the full dioscRi pipeline, and predictive performance was assessed using AUC scores for CAD status. Our dioscRi (AUC = 0.79) outperformed cyCombine (AUC = 0.63) and iMUBAC (AUC = 0.55) (Figure 2K). In combination, these results underscore the strength of dioscRi’s normalization strategy, which outperforms alternatives in downstream subject level classification without requiring explicit batch labels.

### Cell type and marker associations provide complementary biological insights

To assess the interpretability of dioscRi, we first examined the coefficients from the overlapping group LASSO model using different feature sets of predicted cell types from models trained on expert gating. Fitting separate models using only proportions or only marker means provided complementary views: the proportions model identified differential abundance of B cells, pDCs, and CD4+ T cells (Figure 3A), of which only pDCs were present differentially in the original gated data [17] while marker means highlighted expression-level differences within other cell populations, including increased CD192 and CD194 in Tregs (Fig 3B). However, the majority of the differences highlighted in the marker means analysis were not present in the original gated data (Supplementary Figure S2A). For example, CD4+ Treg cells do not express CD14, and additionally CD141+ DCs do not express KLRG1. The combined model revealed some associations consistent with prior findings in coronary artery disease (CAD) [17], with increases in CD4+ Treg expression of CD194, CD45RO and Ki67 (Figure 3C). Conventional CD4+ T cells (Tconvs) were negatively associated with CAD through markers such as CD3, CD4, and CD127 (Figure 3C). This reflects the previously documented changes in Tconv [17], since the expression of CD3, CD4 and CD127 differs significantly between naive T conv (reduced in CAD) and effector memory Tconv (increased in CAD). If Tconv had been divided into two populations (naïve and memory) in the original 11 cell type designation, then additional cell type proportions rather than marker means would likely have been identified by the model. This example illustrates how the marker means can, to some extent, compensate for unbalanced cell type choices in the accuracy of the subject-level prediction, although they require more subsequent analysis to determine the true cell types that differ between CAD+ and CADpatients.

**Figure 3.**
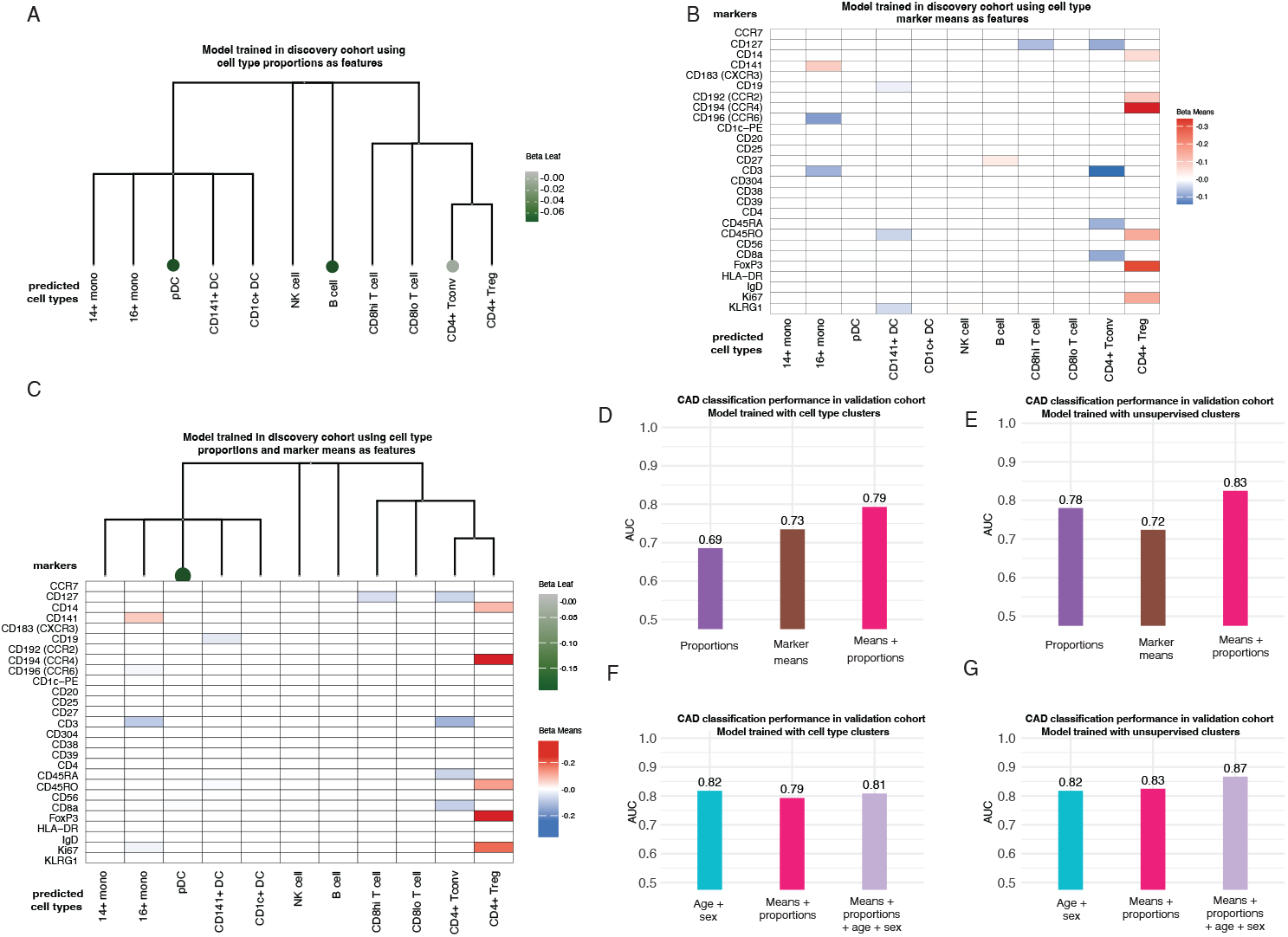
Overlapping group LASSO for the validation cohort identifies cell type associations with CAD status for the predicted cell type model trained using the discovery cohort. (A) Each coefficient corresponds to a predicted cell type proportion: negative values (green) indicate higher abundance in CAD– samples, whereas positive values (purple) indicate enrichment in CAD+ samples. Marker-level coefficients are shown in (B) and combined cell type proportions and marker means in (C): negative values (blue) associate with CAD–, and positive values (red) associate with CAD+. Combining cell type proportions and marker means enhances patient classification for (D) models trained using predicted cell types and (E) models trained using cell type clusters. Incorporating clinical covariates (age and sex) further improves prediction for (F) models trained using predicted cell types and (G) models trained using cell type clusters.

The prediction of CAD from models trained using manually gated cell types is likely limited by the ability of dioscRi to predict the hierarchically defined cell types. Heatmaps of the original gated cell types (Supplementary Figure S3A) and the predicted cell types for the discovery and validation cohorts (Supplementary Figures S3B-C) illustrate these differences. For example, in both the discovery and validation cohorts, expression of CD14, CD183, CD194, CD38, CD39 and CD56 in the three small DC populations is different from that in the manually gated data. The concordance between the size and marker expression of the original manual gated cell types and the predicted cell types is highest for the major cell types such as CD4+ Tconv and CD8hi T cells. Initial choice of 11 gated cell types that were more evenly balanced in size would likely have improved the accuracy of the cell type prediction.

We next trained models that used unsupervised clustering in the discovery cohort to define cell types. When applied to the models built using the unsupervised clusters, dioscRi uncovered additional associations not evident from the annotation-guided analysis (Supplementary Figure S4C). Cluster 7, which had high expression of CD19 and CD20 but also expressed CD1c, CD3, CD4, CD56 and KLRG1 and therefore may comprise a mixture of B cells, DCs and multiple other cell types (Supplementary Figure S4D) showed increased CD25 expression (Supplementary Figure S4B) in CAD+ samples. It is not clear what this represents in biological terms, since cluster 5, whose pattern of marker expression corresponded much better with a pure population of B cells, did not show the same effect of CD25 expression on model performance. Clusters 1 and 4 were characterized by expression of CD56, a marker of NK cells, although their overall pattern of marker expression did not correspond with that of either gated or predicted NK cells (Supplementary Figure S3). Cluster 4 contributes additional information from 3 markers, but the biological relevance is unclear, since NK cells do not express CD14, while expression of KLRG1 is generally low. Thus, while we will next demonstrate that the grouped LASSO approach provides excellent patient-level distinctions, it is difficult to interpret in biological terms when based on preliminary, uncurated unsupervised clustering.

Together, these results illustrate how cell type proportions capture shifts in the abundance of broadly defined populations, while marker means provide finer resolution by revealing differences in cell state within those populations. In this way, dioscRi offers a structured starting point for interpreting complex datasets, using coarse annotations to frame the analysis and marker-level features to uncover additional layers of phenotypic variation, with the understanding that some associations will require further validation.

### Evaluating Predictive Features and Immune Associations

The dioscRi model that is used for patient classification integrates multiple immune features, including cell type proportions and marker means per cell type, which are structured within a biologically or empirically informed hierarchy using a grouped LASSO model. To evaluate the contribution of these components to patient classification performance, we compared three versions of the model: one using only proportions, one using only marker means, and the full model combining both. For models trained using predicted cell types based on manual gates and a biologically derived tree structure from the training cohort, the full model combining proportions and marker means achieved the highest AUC of 0.79 for the validation cohort (Figure 3D) in comparison to the proportions-only model (AUC of 0.69) and the means-only model (AUC of 0.73). As detailed above, the validation cohort data was of poor quality and prediction of cell types without first applying time-gate filtering to remove significant machine artifacts likely compromised the estimation of proportions. Indeed, marker means alone generated a higher AUC than proportions, and inclusion of both further improved model performance. In general, cell type proportions from manually gated data have proven superior in understanding patient-level distinctions. Indeed, in the original published analysis of the BioHEART-CT study [17], there were no significant differences in marker means between the corresponding gated populations in CAD+ and CAD−populations. These results suggest that proportions and marker means provide complementary information that is particularly useful in analysis of datasets that contain substantial technical artifacts.

We next evaluated the patient classification performance for the models that used clustered cell types to annotate the cells. For these models, which also used an empirically derived tree structure, a different pattern emerged. The proportions-only model performed nearly as well as the full model, with AUCs of 0.78 and 0.83, respectively (Figure 3E), while the means-only model had a smaller AUC of 0.72. This stronger performance of the proportions-only model suggests that the clustering process captures biologically meaningful variation in population abundances. These proportions may reflect underlying immune shifts more directly than marker intensities alone, particularly when clusters align well with disease-related immune states. Together, these findings indicate that combining proportions and marker means improves prediction under both annotation strategies.

We further evaluated the added value of dioscRi by comparing its performance to models using only clinical variables. In the original BioHEART-CT study that was designed to describe immune contributions to CAD, independent of known risk factors, SVM models based on cytometry data alone (AUC = 0.65) were inferior to those using only age and sex (AUC = 0.82). In contrast, the normalized clustering-based model in dioscRi is comparable to the age and sex model, achieving an AUC of 0.83 (Figure 3F). Performance improved further when age and sex were included in conjunction with cytometry characteristics, reaching an AUC of 0.87 (Figure 3G). This improvement demonstrates the added predictive value of properly normalized cytometry data, which is especially important for unsupervised clustering.

### dioscRi outperforms state of the art deep learning approaches

We next evaluated the performance of dioscRi in comparison to other deep learning methods, including CellCNN and DeepCNN. We performed this comparison across four cytometry datasets: the BioHEART-CT cohorts, the CMV study SYD519 dataset, the Breast Cancer dataset, and the COVID-19 dataset, each featuring distinct clinical outcomes. For BioHEART-CT, we trained on the discovery cohort and tested on the validation cohort. The COVID-19 and Breast Cancer datasets were split 70–30 into training and testing sets. For the CMV dataset, we followed the split previously described [9], using SYD519 for testing and the remaining studies for training. Across all datasets, we applied a consistent configuration for dioscRi using 11 clusters, while CellCNN and DeepCNN were used with their respective recommended settings. DioscRi outperformed both CellCNN and DeepCNN in all datasets except CMV SDY519 (Figure 4). In the BioHEART-CT validation cohort, dioscRi achieved an AUC of 0.83, well above CellCNN (0.60) and DeepCNN (0.62). Similarly, dioscRi performed strongly on the Breast Cancer dataset (AUC = 0.91 vs. 0.74 and 0.78) and the COVID-19 dataset (AUC = 0.98 vs. 0.70 and 0.57). For the CMV dataset, both dioscRi and DeepCNN achieved high AUCs (0.94), outperforming CellCNN (0.80). These results underscore dioscRi’s robustness and generalizability, particularly in clinical settings where other models show reduced accuracy. In addition to high performance, dioscRi’s interpretable structure facilitates biological insight, a key advantage over more opaque deep learning approaches in which expression of cell markers cannot be readily mapped back to individual cells..

**Figure 4.**
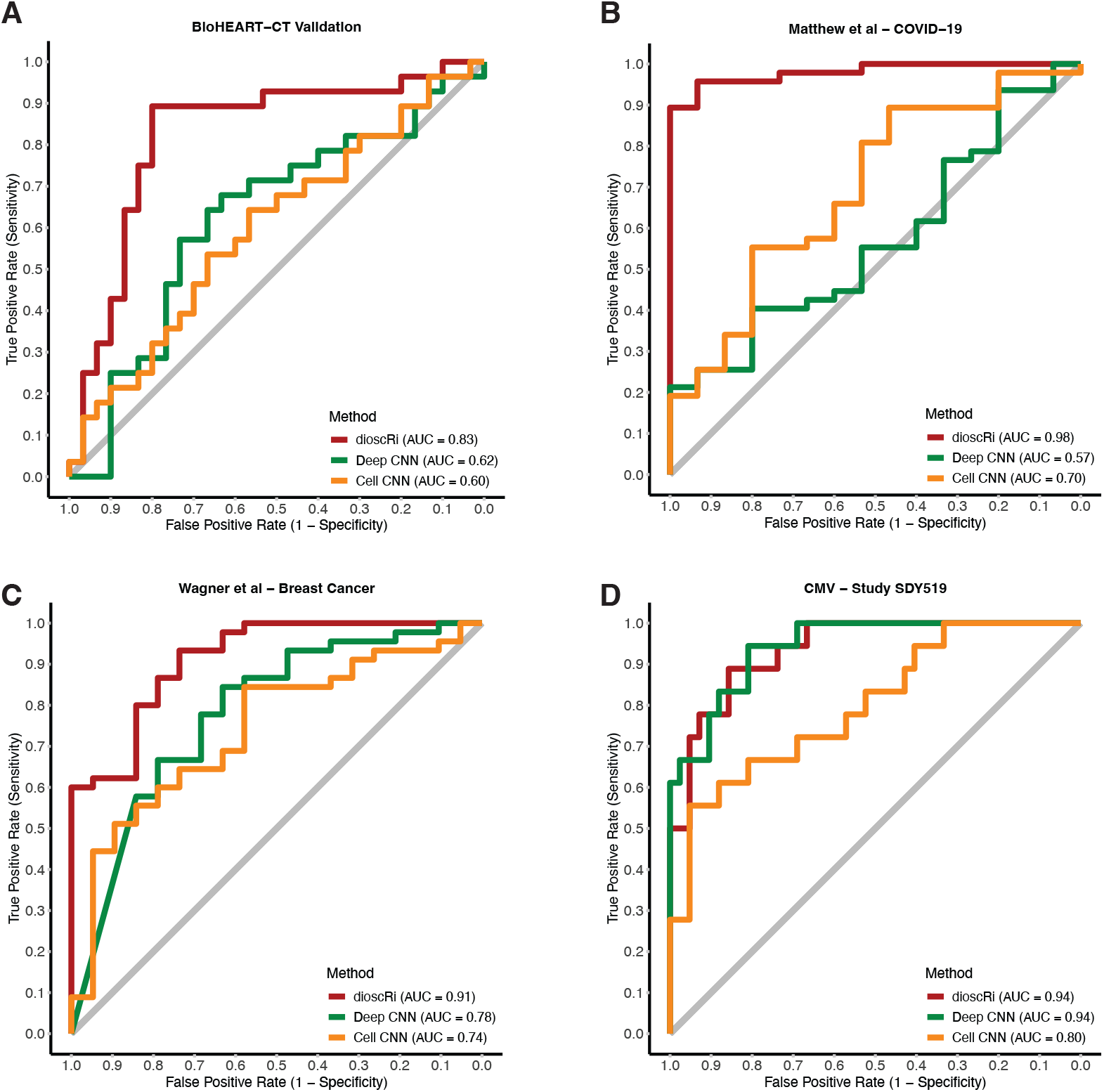
DioscRi outperforms both Deep CNN and Cell CNN across all cohorts, achieving the highest AUCs. When applied to (A) the BioHEART-CT validation study and three external datasets, (B) COVID-19, (C) Breast Cancer, and (D) CMV, the DioscRi normalization-and-prediction pipeline consistently delivers superior classification performance. For the Breast Cancer and COVID-19 datasets, each cohort was split 70% for training and 30% for testing; for the CMV dataset, study SDY519 was reserved entirely for validation.

### Factors Affecting dioscRi’s Performance

The modular nature of dioscRi offers flexibility across datasets, but like many models, its performance can be influenced by several tunable components. We systematically explored the impact of key parameters including the number of clusters, bottleneck layer size, regularization strength (Lambda), and the number of reference samples used for normalization using AUC and total loss as metrics (Supplementary Figure S5). The number of clusters was a strong driver of predictive performance. Across datasets such as BioHEART-CT, CMV, and Matthew et al., we observed that AUC generally peaks at intermediate values, typically around 11 or 12 clusters, before plateauing or slightly declining (Supplementary Figure S5A). This suggests that dioscRi can capture underlying biological structure without overfitting when cluster number is chosen carefully, with cross-validation or heuristic-based approaches likely to help in new datasets. The bottleneck layer size determined how much information was retained in the normalized representation, with moderate sizes (around 15) consistently yielding strong performance (Supplementary Figure S5B). Overly small bottlenecks risked losing critical information, while larger ones tended to overfit. This was reflected in a slight upward trend in validation loss at larger sizes despite total loss remaining relatively stable (Supplementary Figure S5C). Lambda, the regularization parameter in the overlapping group LASSO, shaped the trade-off between model sparsity and performance, with AUC peaking at moderate values (around 0.05) and declining at both lower and higher settings (Supplementary Figure S5D). Total loss remained stable across this range (Supplementary Figure S5E), indicating that tuning Lambda can improve generalization without affecting model stability. Finally, the number of reference samples used in MMD-VAE normalization influenced both performance and consistency. AUC stabilised with around 6 to 7 representative samples, and validation loss improved modestly with larger training subsets (Supplementary Figures S5F-G). These results indicate that dioscRi can achieve strong normalization performance with a relatively small, well-chosen training set, supporting its use even in studies with limited sample sizes. Together, these analyses highlight the adaptability of dioscRi and provide practical guidance for parameter tuning, enabling users to tailor the framework to diverse cytometry applications while maintaining robust predictive performance.

## Discussion

To achieve interpretable and transferable insights from cytometry data, we created dioscRi, a framework combining deep learning normalization with biologically informed feature selection. DioscRi integrates an MMD-VAE for harmonizing multi-batch data and an overlapping group LASSO that leverages cell-type hierarchies to model immune variation. Applied to a coronary artery disease dataset, dioscRi achieved strong predictive performance, and validated a subset of the known associations previously revealed by expert manual gating [17]. Across multiple datasets, dioscRi outperformed existing deep learning approaches while preserving interpretability, demonstrating its value for both accurate prediction and biological discovery.

A major strength of dioscRi lies in its use of hierarchical group structures to model both cell type proportions and marker expression. This allows it to exploit immune system organization to improve interpretability relative to models based on unsupervised clustering. Unlike black-box deep learning models, where feature contributions often require post-hoc explanation tools like SHAP, dioscRi yields coefficients that may represent associations between immune features and clinical outcomes. While the incorporation of biologically defined hierarchies enhances interpretability by maintaining a degree of alignment with known cell type relationships, our results also support previous findings [15] showing that empirically derived hierarchies can be similarly effective. This balance between biological grounding and data-driven flexibility enables dioscRi to provide biological insights without compromising predictive performance.

Our analysis of the BioHEART-CT cohort demonstrates how dioscRi can enhance both prediction and biological insight from cytometry data. In the original BioHEART-CT study [30], SVM models based solely on immune features showed modest predictive performance and did not improve upon simple clinical variables such as age and sex. Using overlapping group LASSO models, we observed more stable and accurate predictions from LDA models trained on either the original manually gated dataset or unsupervised clustering approaches. These improvements were supported by the framework’s normalization strategy and its structured modeling of cell type proportions and marker expression. DioscRi reproduced some previously reported associations, such as decreased pDCs in CAD, and identified marker-level signals that had been obscured by the initial choice of cell types. This included features related to the previously known differences in the population sizes of naïve and memory CD4+ Tconvs. Importantly, integrating cytometry features with the clinical variables age and sex further improved predictive performance, illustrating the added value of immune profiling when supported by effective normalization and feature selection.

The complementary use of predicted cell type proportions and marker means dioscRi captures immune variation at different resolutions. The proportions reflect differences in cell abundance, while the marker means capture changes in phenotype within the predicted cell types. In practice, the line between these two views is shaped by clustering resolution, where higher-resolution clustering can convert marker-level differences into abundance shifts by dividing broader populations into distinct subtypes. Interestingly, the proportion model performed particularly well with unsupervised clusters, likely because the clustering emphasized sample-level heterogeneity in dominant populations. These findings suggest that proportions can serve as strong predictors when clustering effectively captures meaningful immune variation. Although the combined model ultimately performs best, the strong performance of the predicted cell type model highlights its interpretability and value in workflows driven by data-derived cluster structures.

Despite its strengths, dioscRi has some limitations. While the VAE-based normalization reduces technical variability and enables transfer across batches, it does not explicitly model batch differences. In cases with substantial differences, such as shifts in instrument settings or sample handling, residual batch effects may persist and moderately influence downstream analyses. In addition, subtle panel variations, including different antibody clones or minor protocol changes, can introduce technical variability in marker expression that is difficult to distinguish from the biological signal. Although dioscRi’s latent representation is robust to many such differences, these sources of variation could impact performance when applying the model across datasets with substantial experimental heterogeneity. In practice, careful study design and harmonization of upstream protocols can mitigate many of these effects.

In summary, dioscRi offers a robust and interpretable framework for extracting biologically relevant insights from cytometry data and applying them to clinically important prediction tasks. By integrating transferable deep learning–based normalization with structured modeling of cell type features, it overcomes key limitations of existing approaches and improves both performance and interpretability. Across diverse datasets, dioscRi demonstrates consistent predictive strength. These capabilities position dioscRi as a powerful tool for advancing immune profiling in translational research and clinical applications.

## Data availability

Publicly available data were used for all evaluations. All data were downloaded as described in the originating manuscripts.

## Code availability

The *dioscRi* R package is available on Github and is available under the GPL-3 license.

## Acknowledgments

The authors thank all their colleagues, particularly those at Sydney Precision Data Science Centre, and Charles Perkins Centre, for their support and intellectual engagement.

The following sources of funding for each author are gratefully acknowledged: the AIRinnoHK programme of the Innovation and Technology Commission of Hong Kong to JY and EP. Australian Research Council Discovery Early Career Researcher Award (DE200100944) funded by the Australian Government to EP; and the University of Sydney Postgraduate Excellence Award for E.W.; The funding sources had no impact on the study design; in the collection, analysis, and interpretation of data, the writing of the manuscript, and in the decision to submit the manuscript for publication.

## Author contributions

EP and EW conceived and designed the study. EP and EW led the method development and guided the evaluation data analysis with input from HM and JY. BF generated the original gated discovery and validation cohorts that were then curated by EW who implemented all data analytics, and developed the corresponding R code. EP and EW drafted the initial manuscript with writing contributions from SR, and all authors read and reviewed the text. BF extensively edited the manuscript and figures to provide additional clarity related to the immunological aspects of the study. All authors approved the final version.

## Conflict of interest

The authors declare that they have no conflict of interest.

## Supplementary

### Supplementary Tables

### Supplementary Figures

**Supplementary Figure S1.**
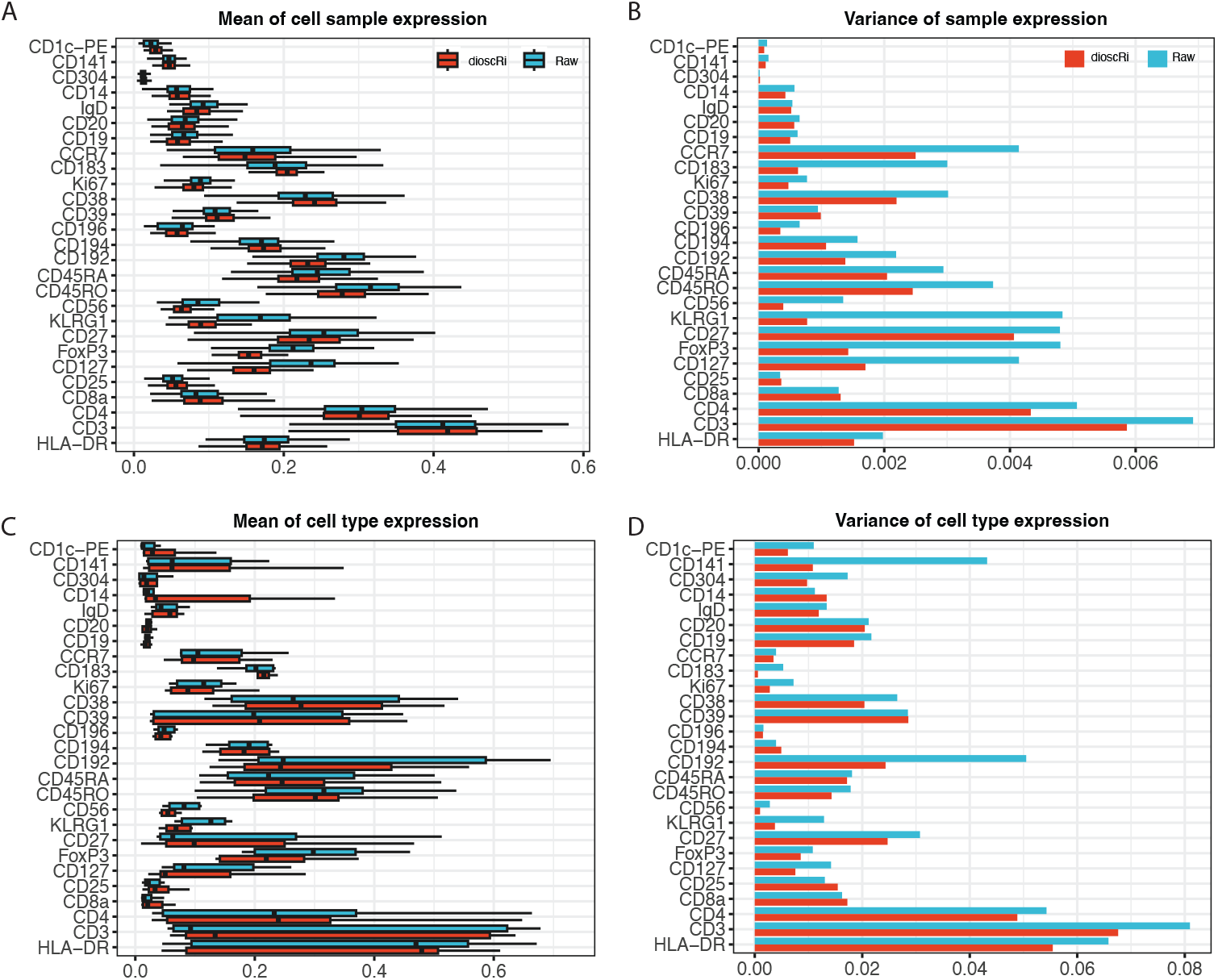
Distribution plots demonstrating the effectiveness of dioscRi normalisation for the discovery cohort using the original manual gates. (A) Each bar shows distribution of the mean expression of the indicated marker within the 111 samples in the discovery cohort. (B) Variance of the means shown in (A). (C) Each bar shows distribution of the mean expression of the indicated marker within the 11 manually gated populations in the discovery cohort. (D) Variance of the means shown in (C)..

**Supplementary Figure S2.**
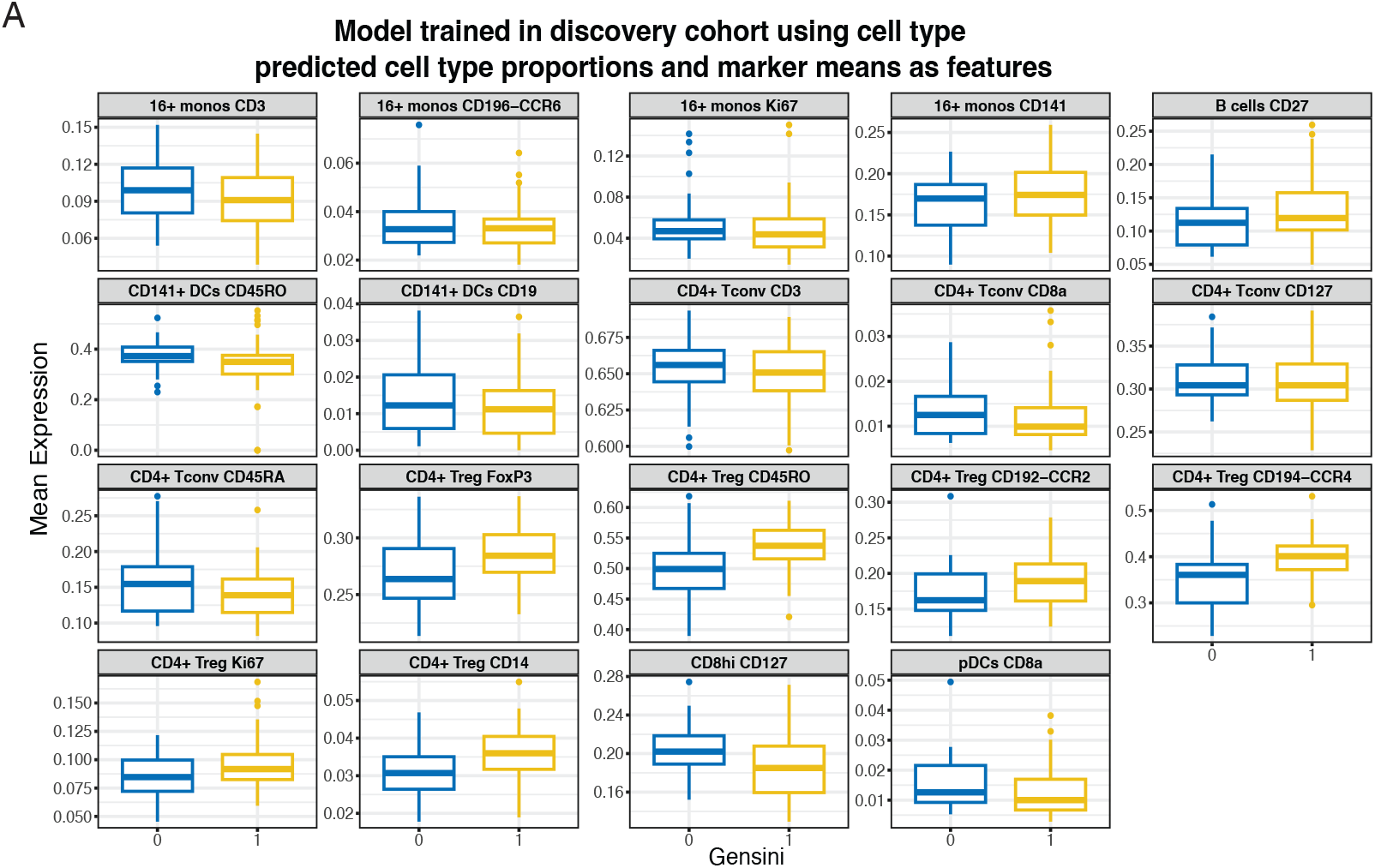
(A) Sample level boxplots of non-zero coefficients of the marker means from the overlapping group lasso model illustrated in Figure 3B-C, split by Gensini (0 = CAD−, 1 = CAD+).

**Supplementary Figure S3.**
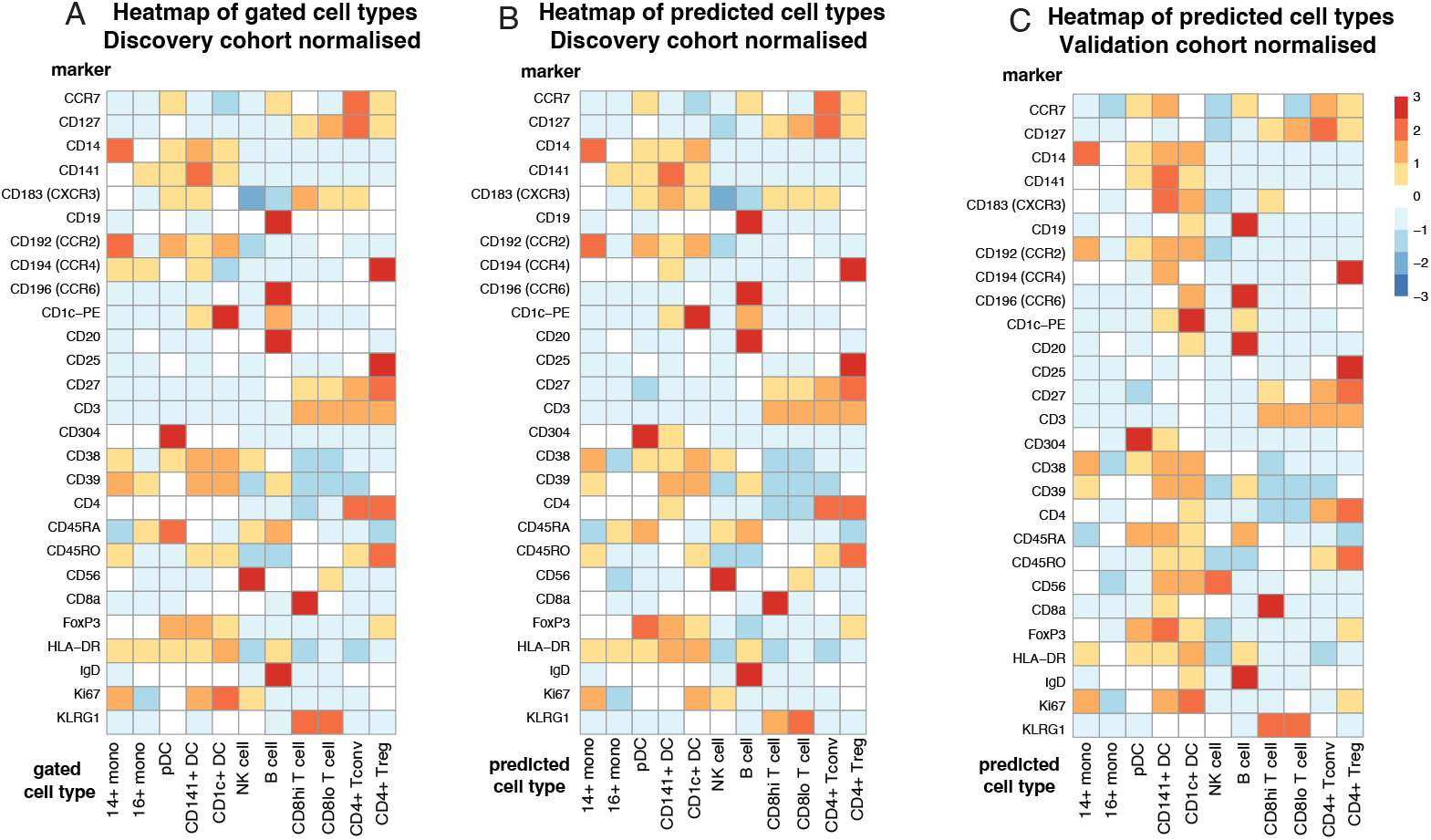
Heatmaps of mean marker expression for normalized data showing 11 cell types from the discovery and validation cohorts. (A) Original manually gated populations, discovery cohort. (B) Predicted cell types, discovery cohort. (C) Predicted cell types, validation cohort.

**Supplementary Figure S4.**
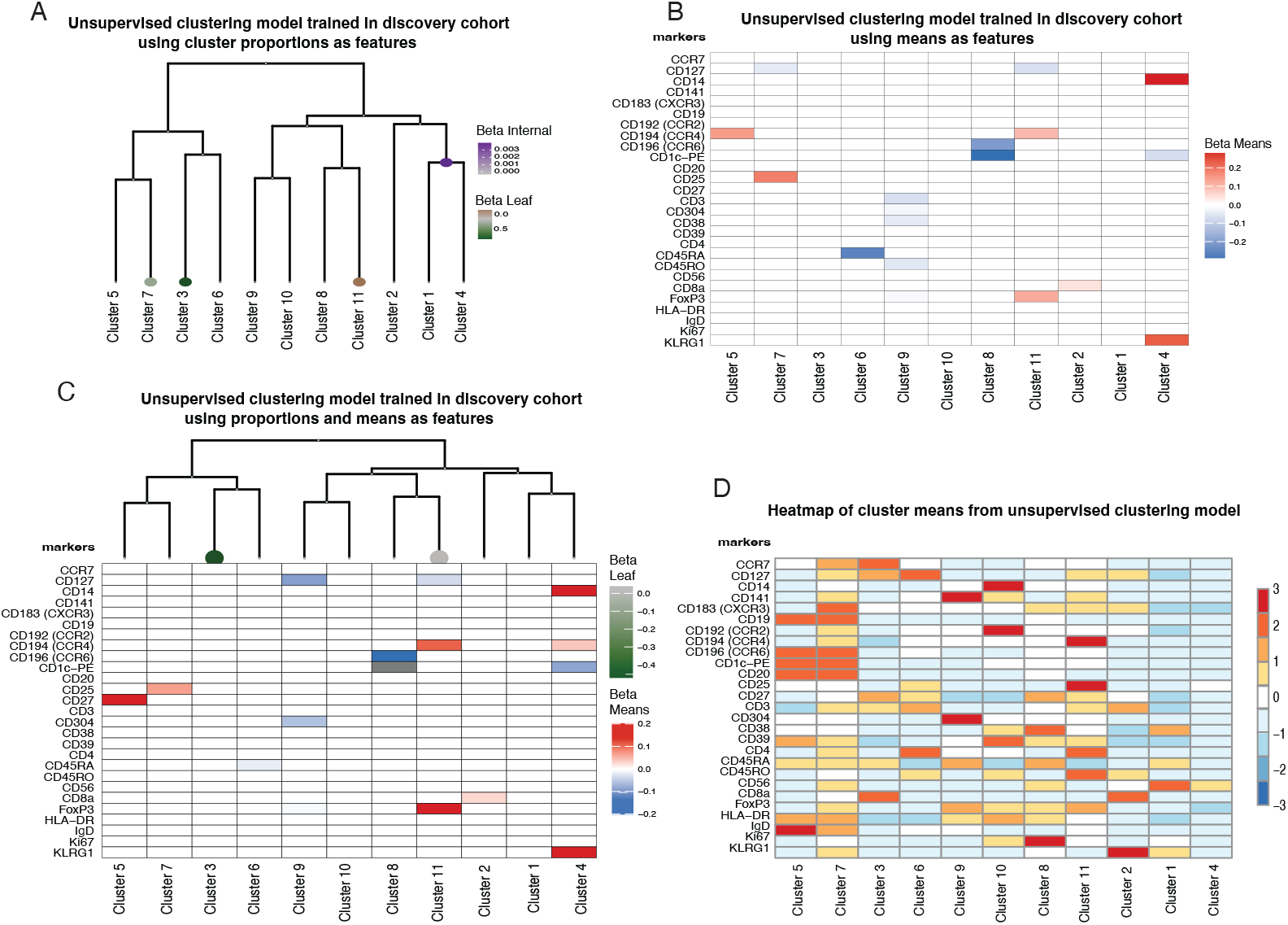
Overlapping group LASSO identifies cell-type associations with CAD status in unsupervised cell clusters. In (A,C), each coefficient corresponds to a cluster proportion: negative values (green) denote higher abundance in CAD– samples, while positive values (purple) denote higher abundance in CAD+ samples. In (B,C), marker level coefficients are shown: negative values (blue) associate with CAD– samples, and positive values (red) associate with CAD+ samples. (D) Heat map of marker expression in clusters.

**Supplementary Figure S5.**
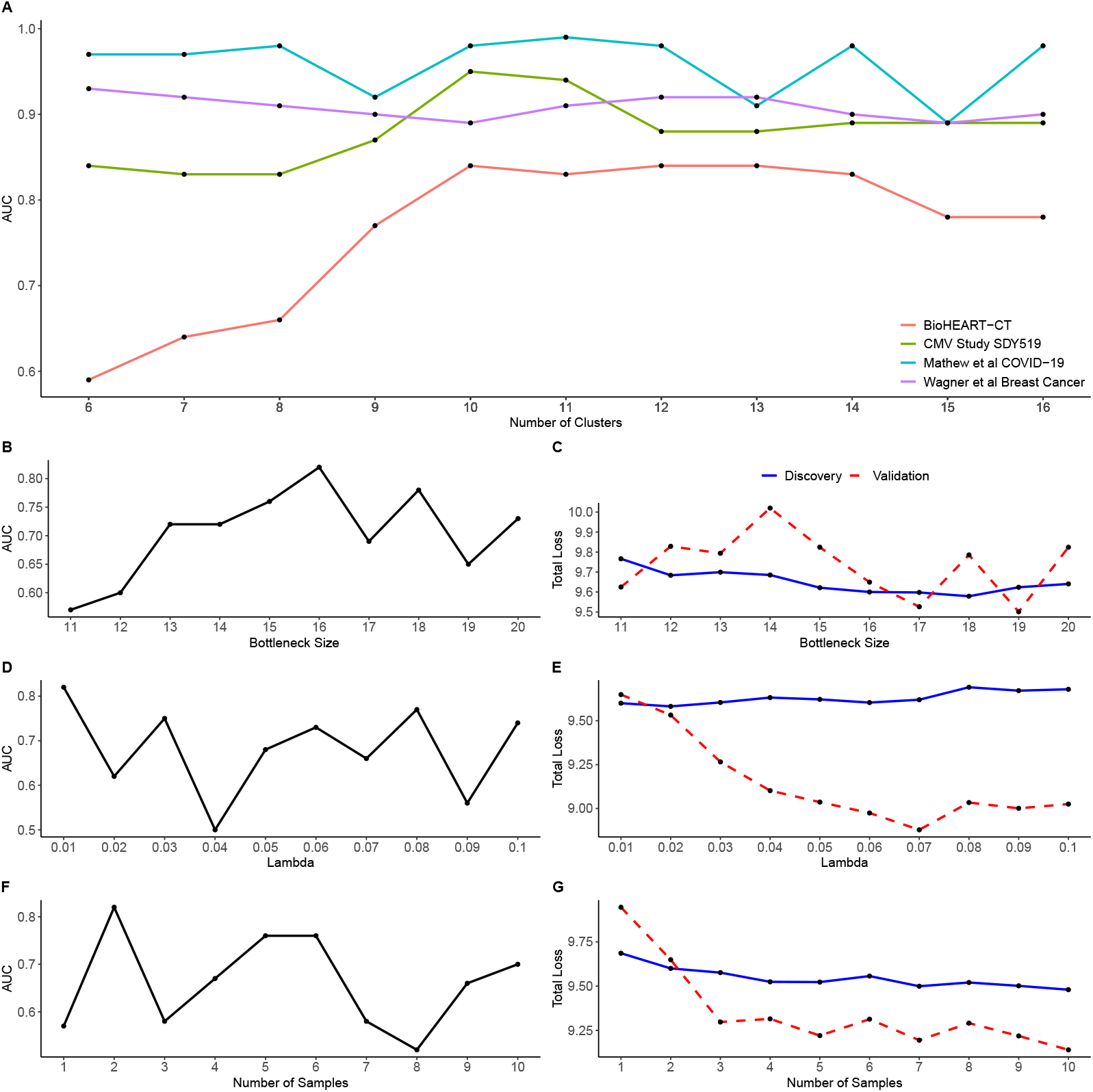
Hyperparameter sweeps identify key parameters affecting dioscRi performance. A) AUC values for CAD status prediction against the number of clusters used during modeling. B) AUC values for CAD status prediction against the bottleneck size for the MMD-VAE normalization. C) Total loss for the discovery and validation studies against the bottleneck size for the MMD-VAE normalization. D) AUC values for CAD status prediction against Lambda for the MMD-VAE normalization. E) Total loss for the discovery and validation studies against Lambda for the MMD-VAE normalization. F) AUC values for CAD status prediction against the number of reference samples for the MMD-VAE normalization. G) Total loss for the discovery and validation studies against the number of reference samples for the MMD-VAE normalization

## Notes

### Competing Interest Statement

The authors have declared no competing interest.

